# A High-Throughput Microphysiological Liver Chip System to Model Direct and Idiosyncratic Drug-Induced Liver Injury Using Human Liver Organoids

**DOI:** 10.1101/2023.12.01.569575

**Authors:** Charles J. Zhang, Sophia R. Meyer, Max A. Garcia, Megan C. Procario, Sanghee Yoo, Amber L. Jolly, Sumin Kim, Jiho Kim, Kyusuk Baek, Roland D. Kersten, Robert J. Fontana, Jonathan Z. Sexton

## Abstract

1.

**Objective:** Drug-induced liver injury (DILI) is a major failure mode in pharmaceutical development. This study aims to address the limitations of existing preclinical models by introducing a high-throughput, microfluidic liver-on-a-chip system, termed as “Curio Barrier Liver Chips,” seeded with human liver organoids to enable metabolic and phenotypic morphologic characterization.

**Methods:** Curio Barrier liver chips, fabricated in an 8×2 well configuration, were utilized to establish 3D liver organoid cultures. Human-induced pluripotent stem cells (iPSCs) were differentiated into hepatic-like organoids (HLOs), and their viability, liver-specific functions, and pharmacological responses were assessed over 28 days.

**Results:** The Curiochips successfully maintained liver physiology and function, showing strong albumin secretion and cytochrome P450 (CYP) activities through 28 days. Unlike traditional models requiring millimolar drug concentrations to detect hepatotoxicity, this platform showed increased sensitivity for APAP and FIAU at micromolar concentrations. *In situ* differentiation of foregut spheroids to liver organoids was also achieved, further simplifying the establishment of liver chips. Furthermore, the chips demonstrated viability, function and DILI responsiveness for 28 days making this an improved model for studying idiosyncratic DILI with prolonged drug exposure and high-throughput capabilities compared to other available systems or primary human hepatocytes.

**Conclusions:** The Curiochips offer an advanced, miniaturized *in vitro* model for early-stage drug development and a sensitive, responsive and cost-effective means to detect direct hepatotoxicity. The high-throughput capability, coupled with robust functionality and pharmacological responses make it a promising tool for improving the prediction and understanding of DILI mechanisms in general and those that required prolonged drug exposure. The model also opens new avenues for research in other chronic liver diseases.

## 2. Introduction

Drug-induced liver injury (DILI) represents a significant challenge in both medical practice and drug development.^1,2^ The liver, being the principal organ for drug metabolism and detoxification, is highly susceptible to damage caused by drug compounds. Most DILI risk is not adequately predicted in available preclinical models or in-vitro test systems, nor observed in phase 1/2 clinical studies. The late-stage detection of DILI for novel molecular entities in clinical studies and post-FDA approval represents a main failure mode in drug development and a leading cause for FDA regulatory actions.^3,4^ Indeed, an estimated 22% of clinical trial failures and 32% of market withdrawals of novel molecular entities are due to unanticipated hepatotoxicity.^5,6^ Although the annual incidence of DILI from all causes in the US general population is estimated at **10 to 20** cases per **100,000** individuals^7^, the incidence of DILI with most drugs is only 1 in 10,000 to 1,000,000 treated patients.^8,9^ Nonetheless, idiosyncratic DILI causes 10% of all acute hepatitis and accounts for nearly half of the cases of acute liver failure in the United States.^10^ Most instances of DILI are termed “idiosyncratic,” (iDILI) since they are largely independent of the dose and duration of drug use and develop in only a small proportion of treated patients (<1 in 1,000). As there are currently over 2,500 prescription medications and >80,000 herbal and dietary supplements available for use in the United States,^11^ the potential for additive or synergistic liver toxicity is high and the currently available *in vivo* and *in vitro* DILI models have low and inadequate predictive capability.^12^

Recent advancements in multicompartment *in vitro* microphysiologic liver-on-a-chip systems that use pumps to achieve tangential media flow have allowed for higher fidelity preclinical prediction of hepatotoxicity compared to conventional 2D monolayer culture systems.^12–14^ While many of these microfluidic models maintain cadaveric primary human hepatocytes (PHH) or iPSC-derived liver organoids for longer times than traditional assays, there are still prominent differences in culture longevity, cost, throughput, and scalability with existing platforms.

The lack of viable long-term in vitro liver models poses a significant limitation in the capacity to investigate both direct and idiosyncratic DILI. In most instances, a direct hepatocellular toxicity response is observed that leads to immediate liver damage. However, clinical evidence indicates that DILI often manifests weeks or even months into a treatment regimen, or even weeks after the discontinuation of a drug.^15,16^ Clinically, idiosyncratic DILI also exhibits a high degree of heterogeneity in its presentation and severity. In many cases, liver injury does not improve following the withdrawal of the offending drugs, and it may even continue to worsen. Our understanding of the genetic and environmental factors contributing to idiosyncratic DILI risk remains limited despite extensive in vitro and preclinical testing of all new molecular entities during their development.^17,18^ These complex phenomena cannot be effectively modeled without the availability of long-term hepatocyte culture models that maintain physiological relevance for extended periods after seeding or patterning with multiple liver and immune cell types. In addition, the demand for an effective long-term liver model extends beyond the study of idiosyncratic DILI with prolonged drug dosing. Other common chronic liver diseases such as metabolic-associated steatotic liver disease (MASLD), fibrosis, and chronic viral hepatitis also highlight the unmet need for robust, long-term multi-cellular liver models to facilitate future natural history and mechanistic studies.

This study describes the development of a high throughput, robust, and low-cost microphysiological 3D platform for drug hepatotoxicity testing that relies on passive osmotically-driven media flow, in contrast to existing chips that use pumps to circulate media. In this study, we demonstrate the functionality of iPSC-derived multicellular human liver organoids (HLOs) on the Qureator Curio Chip platform along with the phenotypic and metabolic functionality of the hepatocytes in culture media followed by experiments showing their metabolic activity and ability to serve as models for necrotic (APAP) and mitochondrial (FIAU) mechanisms of direct hepatotoxicity. Finally, we demonstrate the ability of this microphysiologic system to maintain cellular health and function for up to 28 days which will allow for experiments involving prolonged drug dosing implicated in idiosyncratic DILI.

## 3. Results

### 3.1 Design and Engineering of a High-Throughput Microfluidic Liver-Chip

Curio-Barrier chips, developed by Qureator Inc., were engineered with 16 wells in an 8×2 configuration allowing for the establishment of 3D liver organoid cultures embedded in a mixture of collagen type 1 and Matrigel. Wells were designed to allow for patterning in three channels (side 1, side 2, and the middle channel), allowing for varying hydrogel and cell types without membrane separation and including two media reservoirs. Curio-Barrier chips are SBS plate format compatible to facilitate HTS screening using a liquid handler and high-content imaging-based characterization. The 3D liver organoids were dispensed in the middle channel of the Curiochip and an acellular hydrogel was pattered into the side 1 channel. The culture medium was dispensed into the side 2 channel (Figure 1).

**Figure 1.**
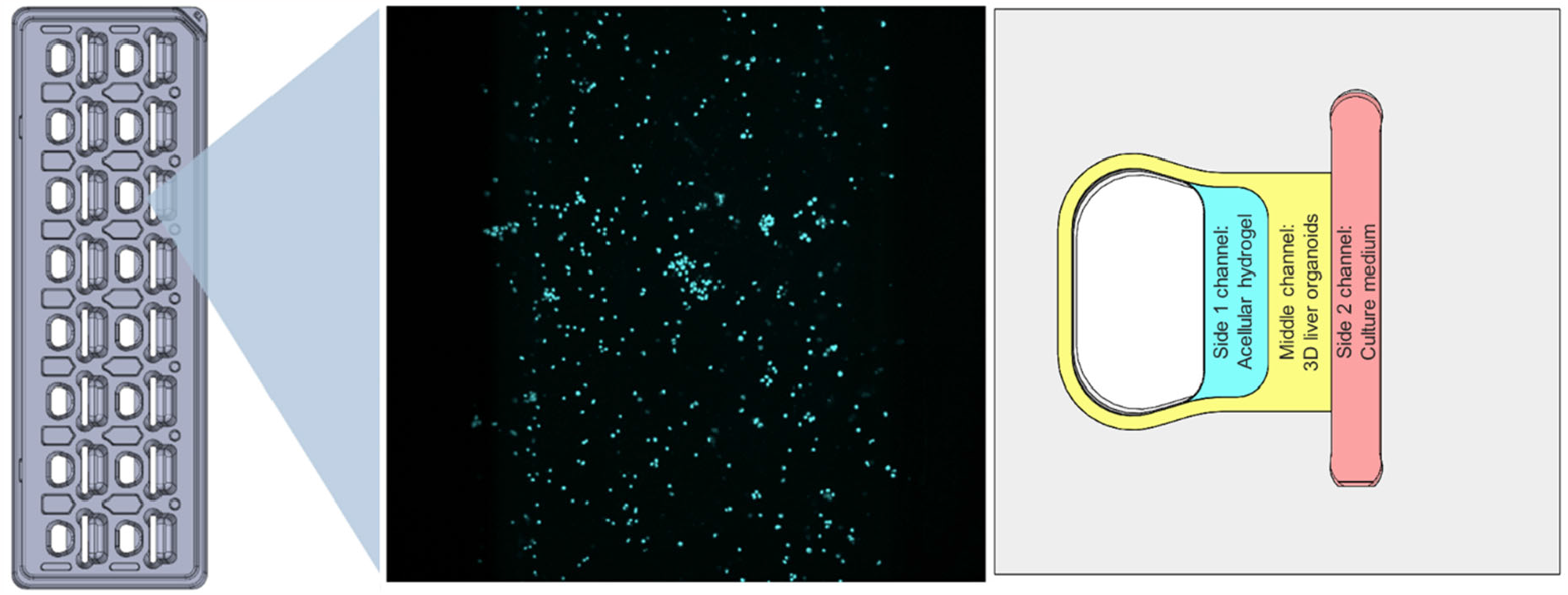
Diagram of a 16-well HLO Curiochip with fluorescent image of a single well and detailed configuration.

### 3.2 Maintenance of Hepatocyte Viability and Metabolism in High-Throughput Curiochips for up to 28 Days

For these studies, human iPSC line 72.3 cells obtained from Cincinnati Children’s Hospital Medical Center were differentiated into HLOs based on a previously described protocol.^14,19^ Each well contained two media ports and for our purposes of developing HLO liver chips, these ports were used with media gradients to achieve a steady hydrostatic media flow. With an emphasis on designing a high-throughput, SBS-plate compatible format, media collection, and maintenance was achieved in Curiochips by standard multichannel pipettors/automated liquid handlers.

Dispersed HLO cells dispensed and cultured in Curiochips with hydrostatic media flow continue to proliferate and retain hepatocyte and stellate markers across 7 days of culture (Figure S1). Visually, hepatocytes maintained a rounded morphology and continued to grow in clusters whereas hepatic stellate cells exhibited an extended fibrotic phenotype. Curiochips show a consistent increase in albumin expression across a 7-day culture period, comparable to our previous reports of albumin secretion by hepatocyte-like cells on an established microfluidic chip system (Figure 2A).^14^ Curiochips on day 7 show CYP 1A, 2D, and 3A family activities based on metabolic turnover of acetaminophen, cyclophosphamide, and darunavir, respectively (Figure 2C). While CYP 1A and 3A family activities are comparable to previously developed liver chips, metabolic turnover is less than from liver chips seeded with validated lots of PHH. However, HLOs consist of 65-70% hepatocytes, and when scaled for the total number of hepatocytes, albumin secretion is nearly comparable to PHHs (Figure S2) indicating enhanced liver function. CYP 2D showed a significant increase in expression compared to previous HLO microfluidic liver chips, nearing the enzymatic activity of PHH liver chips. In addition, after 24 hours of incubation with drugs, increased expression of CYPs 1A2, 2D6, and 3A4 is observed, suggesting a degree of CYP induction (Figure 2B). Furthermore, absorption studies with Curiochips patterned with hydrogel show no inherent compound absorption by the device (Figure S3).

**Figure 2.**
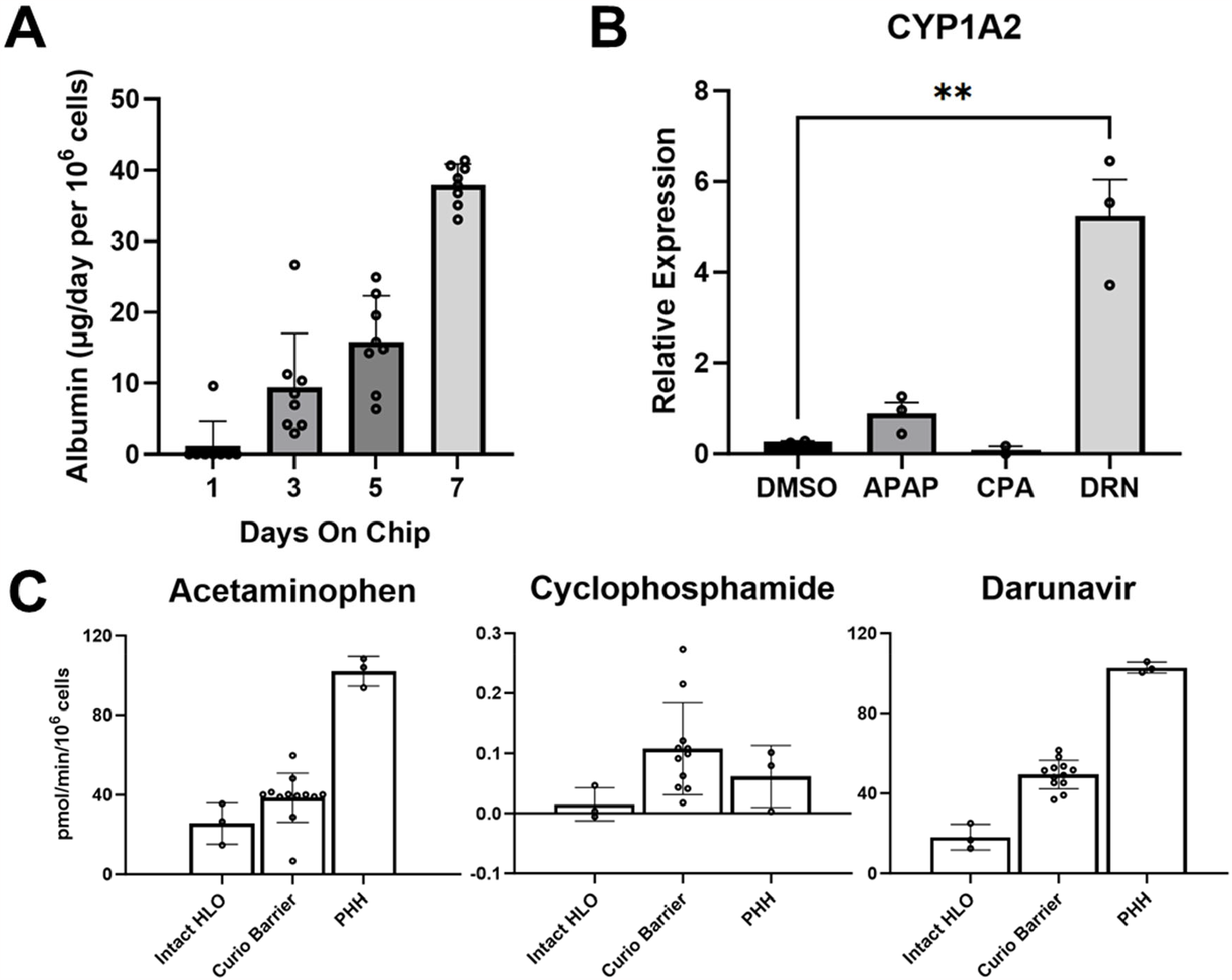
Dispersed HLO growth on Curiochips show A) continuous increase in albumin production across 7 days of culture, B) induction of CYP1A2 by acetaminophen (APAP) and darunavir (DRN), and C) increased CYP450 activity of HLO Curiochips compared to intact embedded HLOs.

### 3.3 28-Day Microfluidic Liver Chips as a Non-Acute/Chronic Model of Idiosyncratic DILI

While our findings in HLO adaptation to Curiochips matched previous findings in other microfluidic systems, we believe that either space restrictions or lack of hydrogel embedding were limitations in the long-term maintenance of liver chips to model idiosyncratic DILI in lower-dosage and prolonged treatment regimes. To this end, Curiochips were assessed for albumin secretion and CYP activity over 28 days. HLO cells were plated at days 0, 7, 14, and 21 and were monitored for albumin secretion and CYP activity for 7 days at each time point. Periodic assaying of output media demonstrated consistent albumin production through day 21 with reduced but persistent secretion measured at 28 days. CYP expression was variable week-to-week, but significant diminishments were not identified until day 28 (Figure 3A).

**Figure 3.**
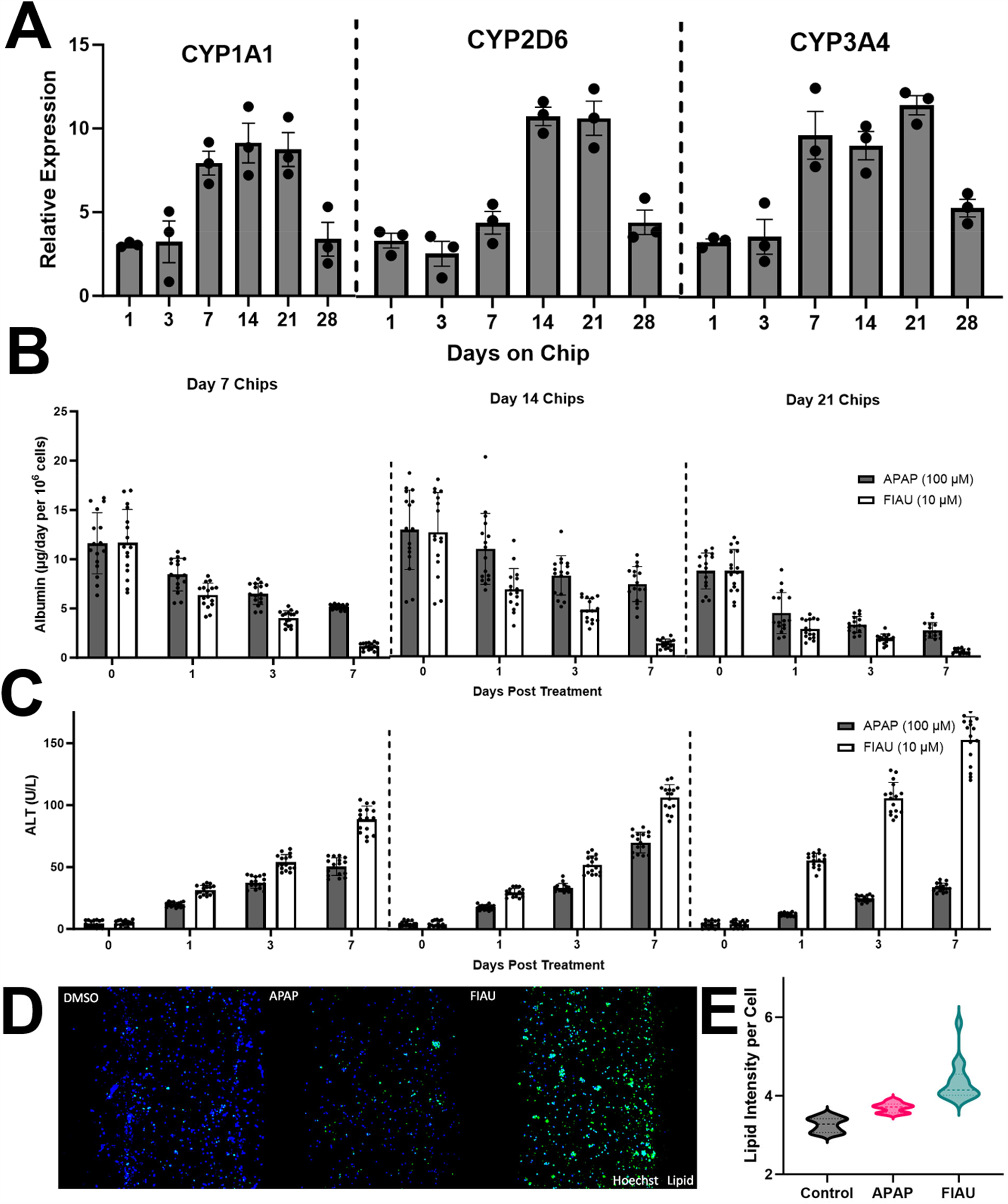
Establishing a long-term liver chip for DILI modelling. A) CYP450 levels increase through day 7 and are maintained through day 21. Day 7, 14, and 21 HLO Curiochips are viable models for assessing APAP and FIAU-induced DILI in both B) albumin and C) ALT responses. D) Fluorescence imaging of Curiochips show increased lipid accumulation in APAP or FIAU treated chip wells where blue is Hoechst-33342 and green is HCS LipidTox Green. E) Quantitation of lipid accumulation at the single cell-level measured using Cell Profiler.

Strong albumin secretion was observed weekly across 28 days with HLOs on the Curiochip platform (Figure 3B). Curiochips at days 7, 14, and 21 were compared for DILI response in terms of ALT release into the media compartment to assess the model viability across time using control DILI drugs acetaminophen (APAP) and fialuridine (FIAU). Increased ALT activity in collected media is observed from both APAP and FIAU treatments with 7 days of drug treatment across chips treated after 7, 14, and 21 days in culture (Figure 3C). Liver chips show hepatocellular injury through increased ALT in response to APAP at 100μM, suggesting adequate CYP metabolism to the toxic metabolite N-acetyl-p-benzoquinone imine (NAPQI). Conventional DILI assessment in static cultures with PHH monolayer sandwich cultures or 3D spheroids generally requires 1-10mM APAP concentrations to produce significant hepatocellular injury.^20,21^ A distinct advantage of this system is the observed enhanced potency of APAP resulting in hepatocellular injury that is comparable to hepatotoxic serum APAP concentrations in humans (1 microgram/mL or 6.6μM APAP).^22^ FIAU also produced a large increase in ALT release at 10μM (Figure 3B and 3C), consistent with our previous studies.^22^ In addition, measured total albumin decreased after treatment with FIAU and APAP with low well-to-well variability. Interestingly, day 7 and day 14 liver chips demonstrated identical response patterns to APAP and FIAU in both albumin production and ALT release throughout an additional 7 days of culture with drug treatment. Day 14 liver chips demonstrated a similar pattern of staining for APAP and FIAU with marked changes in lipid content (Figure 3D). At day 21, liver chips showed a unique hepatotoxic response where significantly higher ALT release was observed following FIAU treatment suggesting worsened cell health at this timepoint. Inversely, APAP-treated liver chips showed minimal ALT release. As we observed a reduction in CYP expression nearing 28 days of culture, we predict that this reduction in hepatocellular injury is a result of reduced metabolic conversion of APAP to NAPQI but confirmatory metabolomic studies are needed.

After media was sampled for biochemical/metabolomic analyses, liver chips were then fixed and stained for hepatocyte and stellate markers at days 7, 14, and 21 and showed comparable marker expression in both percent positivity and fluorescent intensity (Figure 4A). The chips were stained with Hoechst-33342 and HCS LipidTox Green and imaged using high-content confocal microscopy to gain insight into the mechanism of DILI by examining the phenotypic presentation of HLO cells. APAP treatment appeared to result in necrosis as observed by an overall reduction in cell counts, indicative of hepatocyte death (Figure 3D). Treatment with FIAU showed a characteristic accumulation of neutral lipid droplets (Figure 3E), mirroring the observed histologic presentation of steatohepatitis with FIAU from mitochondrial toxicity.^23^

**Figure 4.**
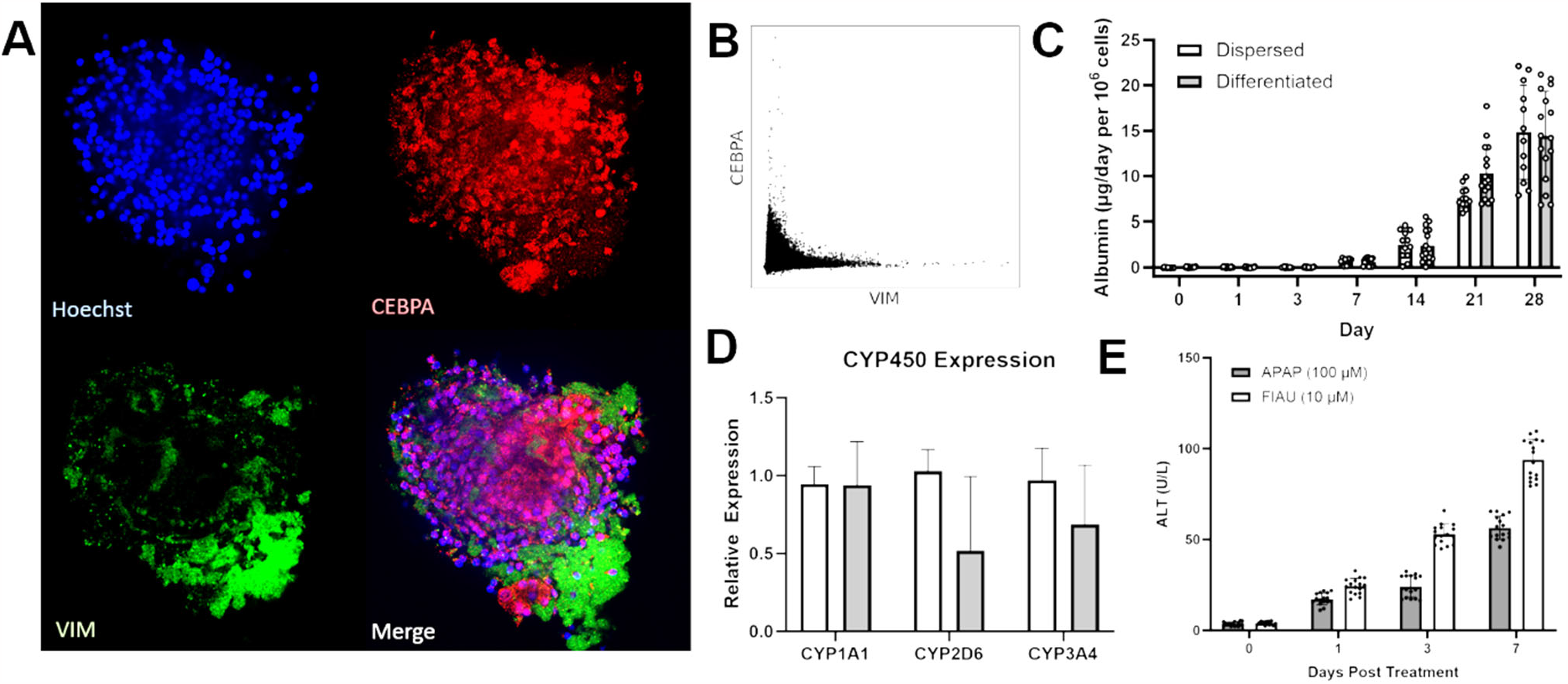
Differentiation of HLOs on Curiochips. A) Confocal fluorescence imaging of HLOs on Curiochips grow as liver spheroids and show cell-specific staining of hepatocyte (red – CEBPA) and stellate markers (green – vimentin). C) Albumin production of HLOs differentiated on Curiochips match that of cells dispersed onto chip. D) CYP450 expression is consistent across cells dispersed into Curiochips and HLOs differentiated on chip. E) HLOs differentiated on chip exhibit response to DILI compounds APAP and FIAU.

#### In Situ Differentiation of Foregut Spheroids to HLOs on Curiochips

As the Curiochips allow for dispensing HLO cells embedded in hydrogel, we sought to test the capability of an *in situ* differentiation of definitive endoderm spheroids to HLOs within the device. Upon full completion of the standard protocol, organoids on liver chips showed positivity for hepatocyte marker CEBPA, HNF4A, HIF1A, and ASGR1, while also positive for stellate markers vimentin, reelin, and α-SMA (Figure 4A, CEBPA and vimentin are shown). High-content imaging demonstrated that hepatocyte and stellate marker staining is regional, showing selective liver cell differentiation at percentages comparable to off-chip differentiation (Figure 4B) of HLOs cultured in standard Matrigel embeddings until day 21.

Media outflow shows substantial albumin secretion as early as 7 days post hepatocyte growth media (HGM) treatment with a gradual increase through 14 days of treatment (Figure 4C). Interestingly, while albumin levels match that of dispersed HLOs transferred to Curiochips at an earlier culture date, we observed no evidence of reduced function by any metric. First, peak albumin expression was comparable between native differentiated and transferred liver chips. Also, CYP450 expression and metabolic turnover showed no significant changes for any CYP family substrates, nor were there any observations of changes to CYP induction (Figure 4D). Lastly, native liver chips show similar ALT response patterns to APAP and FIAU (Figure 4E) along with identical morphological perturbations in lipid staining (Figure 3E) despite differing distinct morphologic perturbations. These observations suggest that direct seeding and in situ differentiation of definitive endoderm spheroids may be a reliable model for drug hepatotoxicity testing and further reduce the time, steps, and labor required for testing and associated costs.

## 4. Discussion

Recent efforts have demonstrated the superiority of advanced patient-derived models such as HLOs and their potential for standardization in drug discovery. However, there are limitations to accessibility, replicability, and scalability. Liver chips built around the Curiochip system were designed to mitigate these issues. The iPSC differentiation to HLO protocol was originally designed around existing 2D monolayer culture methods and has recently seen popular adoption in liver research. This contrasts with existing PHH methods which are costly and suffer from substantial batch-to-batch variability and limited availability. Additionally, PHH-based systems are limited in their physiologic relevance due to the inability to co-culture with non-parenchymal cells from the same patient lineage.

Currently available microfluidic liver models present several limitations related to scalability, reproducibility, and consistency. The inherent characteristics of existing microfluidic systems include high costs, poor reproducibility across patient donor cells, and complex drug pharmacokinetics on liver chips. These factors significantly restrict the batch sizes that can be employed, limit the ability to generate replicates, hinder large-scale testing of drugs over a wide dose range, and impede assessments at multiple time points during experiments. Another issue with many microfluidic devices is the choice of materials, often relying on polydimethylsiloxane (PDMS) or other highly drug-adsorbent materials.^24^ These materials can lead to artifactual observations in drug effects and kinetics, potentially compromising the reliability of experimental results. To address these multiple challenges, we have developed an improved scalable alternative that uses iPSC-derived human liver organoids (HLOs) and incorporates them into an engineered high-throughput microfluidic platform, with a primary focus on enhancing consistency across replicates and minimizing inherent drug interactions to ensure robustness, particularly when applied to large-scale compound testing. Additionally, this approach incorporates multiple liver cell types including hepatocytes, stellate, and Kupffer cells that may enhance the predictive power in DILI risk assessment. This system has been designed to integrate with standard laboratory automation, facilitating a wide array of analytical techniques such as bioassays, immunofluorescence, transcriptomics, high-content confocal fluorescence imaging, and image cytometry. Despite its compact size, this liver-on-a-chip architecture retains essential features, including the ability to control media flow with liver organoids embedded into extracellular matrix scaffolds.

We envision that this system can significantly enhance the rigor and reproducibility in pre-clinical DILI risk assessment, thereby reducing attrition in drug development due to DILI and enhancing patient safety in clinical studies and post-FDA approval. Another major improvement enabled by higher throughput methods is the screening of multiple patient-derived organoids to predict the risk of liver injury across patients with diverse genetics who have experienced DILI. Furthermore, the Curiochip sandwich architecture makes it well-suited for testing compounds with longer drug exposure periods, which can be invaluable in detecting DILI in the early stages of preclinical drug development to help prioritize molecular scaffolds for lead optimization and clinical candidate selection.

Although the level of complexity associated with multicellular, 3D cultures, and dynamic fluid flow has shown to be a hallmark in establishing improved models, these features have not been compatible with high-throughput platforms and automated liquid handlers. However, our efforts effectively miniaturize the complex organ-on-chip systems to achieve higher throughput without sacrificing environmental conditions that deliver high physiological performance. In effect, this enables HLO Curiochips to be a cost-effective and rapid means for early-stage drug discovery testing for both efficacy and safety testing. The scalability to 96 wells per plate also allows for more complex, large dose range and multi-drug combination testing compared to larger format microfluidic liver chip systems.

Our findings also suggest that miniaturized liver chips are capable of supporting the long-term maintenance of iPSC-derived liver cells. Currently, models for idiosyncratic DILI with prolonged drug exposure and other chronic liver diseases are limited in terms of the treatment timeframe. HLO Curiochips showed the ability to maintain viable liver cells for up to 28 days and demonstrate their usability in low-dose, long-term studies, or liver recovery/regeneration research. This long-term model may also be useful to study viral hepatitis. Future studies with HLO Curiochips will focus on model development for idiosyncratic DILI using patient-derived HLOs that incorporate inflammatory stressors or same-patient immune cells in co-culture with HLOs.

## 5. Methods

### 5.1 Curiochip Preparation for 3D patterning

A detailed protocol for Curiochip preparation for 3D patterning is described in Kim, et al, 2023.

### 5.2 Human liver organoid differentiation and chip culture

HLOs were differentiated as previously described.^14^ In brief, human iPSC line 72.3 obtained from Cincinnati Children’s Hospital Medical Center was differentiated into HLOs based on a previously described protocol.^19^ iPSCs were seeded at high density on growth factor reduced Matrigel (Corning, 354230) coated 6-well plates (Thermo Fisher Scientific, 140675) to achieve 90% confluency after 24 hours of culture. iPSCs were then treated with Activin A (R&D Biosystems, 338-AC) for 3 days and FGF4 (purified in-house) for 3 additional days to bud definitive endoderm spheroids.

Spheroids were embedded in 75 μL Matrigel (Corning, 354234) droplets in 6 well plates and treated with retinoic acid for 4 days followed by hepatocyte growth media (Hepatocyte Culture Medium BulletKit (Lonza, CC-3198) supplemented with 10 ng/mL hepatocyte growth factor (PeproTech, 100-39), 20 ng/mL oncostatin M (R&D Systems, 295OM050), and 0.1 μM dexamethasone (Millipore Sigma, D4902)) for 12 days. Following culture, cells were removed from Matrigel embeddings by mechanical dislodging with 10 mL wash media (DMEM/F12 supplemented with 1X pen/strep). In a 15 mL tube, organoids in Matrigel were broken apart by repetitive pipetting and spun down at 300 x g for 3 minutes. Media and empty Matrigel were carefully aspirated before repeating washing with fresh 10 mL of wash media. This procedure was repeated until most Matrigel residue was visually removed following roughly 3-5 washes.

Free organoids were then dispersed by resuspension in 0.25% Trypsin and were transferred to a 6-well plate for 37 °C incubation. After 10 minutes of incubation, cells were pipetted up and down and returned to the incubator for an additional 5 minutes. Trypsin activity was stopped by the addition of 1 mL FBS, transferred to a fresh 15 mL tube, and spun down at 500 x g for 5 minutes. Trypsin with FBS was then aspirated and cells were resuspended in PBS and passed through a 100 μm filter. Cells in PBS were then counted for downstream applications.

### 5.3 Curiochip Patterning and Culture

Dispersed HLOs were resuspended in a hydrogel mixture of Rat Tail Collagen Type 1 (Corning, 354236) and Matrigel (Corning, 354230) at a density of 10,000 cells per 1.4 uL and kept on ice. 1.4 μL of this hydrogel cell suspension was pipetted into the middle channel of each Curiochip well and was incubated at 37 °C for 15 minutes to solidify. An additional volume of cell-free hydrogel mixture was loaded into the side 1 channel and the culture medium was patterned into the side 2 channel and incubated at 37 °C for 15 minutes to solidify. 100 μL of complete HGM media was added to the top reservoir while 20 μL was added to the bottom reservoir to create a volume gradient that would allow for gradual flow. Media was collected and changed 1 day after patterning followed by every other day.

For assessing the differentiation of definitive endoderm to HLOs, foregut spheroids from day 7 of the differentiation protocol were resuspended in the hydrogel mixture and loaded into the middle channel of the Curiochips instead of dispersed HLOs. Following solidification and hydrogel addition to the side 1 and 2 channels, the differentiation medias were identical to standard HLO differentiation. 100 μL of each media was added to both reservoirs (no flow) until the addition of HGM, where the 100 μL of media was added to the top reservoir and 20 μL in the bottom reservoir to introduce flow.

### 5.4 Human Serum Albumin and ALT Measurements

50 μL of media from both reservoirs for each chip well was collected and pooled at respective time points. Albumin from chip media outflow was measured using ELISA (R&D Systems, DY1455). 10 uL of media was diluted 100-fold in PBS before incubation on an ELISA plate. A standard curve was created with albumin concentrations ranging from 2.5 to 160 ng/mL. Each sample was assayed in triplicate and results were scaled per 10^6^ cells.

For measuring ALT and AST, 30 uL of media and PBS were dispensed into a 96-well assay plate. 300 uL of room temperature ALT/GPT reagent (Thermo Fisher Scientific, TR71121) or AST/GOT reagent (Thermo Fisher Scientific, TR70121) was then dispensed into all wells in the plate and incubated at 37 °C for 30 seconds before recording absorbance at 340 nm for 3 minutes. The activity of ALT or AST was determined using the following equation:

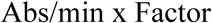

The Factor constant is pre-determined for this assay as 1746, using the manufacturer’s recommendation. The average absorbance from blanks was subtracted from all other samples. Plates were read with a BioTek Synergy H1 Microplate Reader.

### 3.5 CYP450 Mediated Substrate Metabolism

Acetaminophen, cyclophosphamide, and darunavir were chosen as substrates for CYP 1, 2, and 3 families respectively. The media in Curiochips was replaced with 100 μL media in each reservoir containing substrates at 10 μM. After 1 hour and 2 hours of incubation, 50 μL of media from each reservoir was pooled together to achieve 100 μL of total media for analysis. Reactions were stopped using 100 μL of cold methanol and centrifuged for 5 minutes at 3,000 RPM to collect the supernatant. Samples were analyzed by quantitative LC/MS/MS to detect the diminution in the drug parent mass using a Thermo ESI-Q-Exactive Orbitrap mass spectrometer coupled to a Thermo Vanquish ultra-HPLC system. Experimental variables include a Phenomenex Kinetex 2.6 um C18 reverse phase 100 Å 150×3mm LC column, 2 μL injection volume; LC gradient, solvent A, 5% acetonitrile and 0.1% formic acid; solvent B, 100% acetonitrile and 0.1% formic acid; 0 min, 0% B; 1 min, 0%B; 7 min, 80% B; flow rate 0.4mL/min; MS, and positive ion mode was used.

### 3.6 Image Acquisition

Confocal fluorescence images were acquired with a Yokogawa CQ1 or CellVoyager 8000 High-Content Screening System in a custom-fabricated SBS plate format Curiochip holder. Each holder accommodates three Curiochips, totaling 48 wells per imaging cycle. 3D images were acquired on the CQ1 using a 10X/0.7NA objective lens by obtaining 10 Z-planes across a 100 μm range. Imaging on the CellVoyager 8000 system was performed using a 20X/1.0NA water immersion objective lens where 1 μm Z-stacks were obtained across 150 μm to produce maximum intensity projection images for analysis.

## Abbreviations

DILI: drug-induced liver injury
HLO: human liver organoid
PHH: primary human hepatocyte
iPSC: induced pluripotent stem cell
APAP: acetaminophen
FIAU: fialuridine
ALT: alanine aminotransferase

