## Supplemental Figures for "A High-Throughput Microphysiological Liver Chip System to Model Direct and Idiosyncratic Drug-Induced Liver Injury Using Human Liver Organoids"

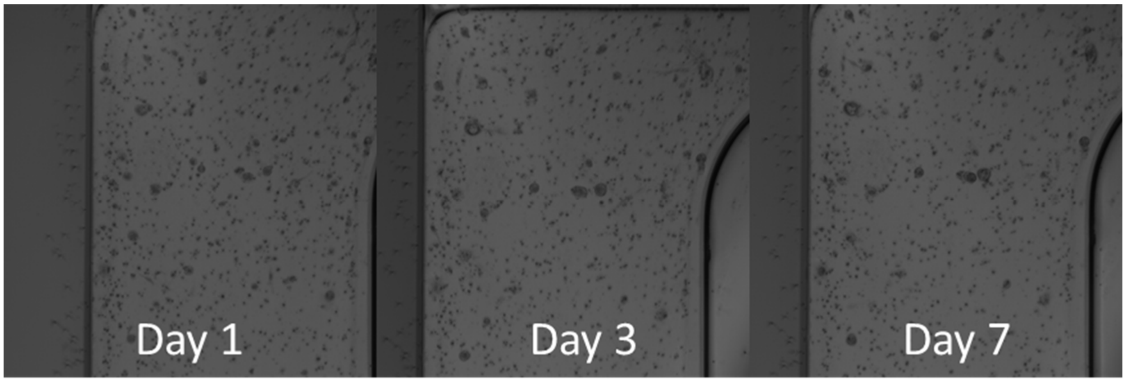


Figure S1. Dispersed HLOs continue to grow when cultured on Curiochips.

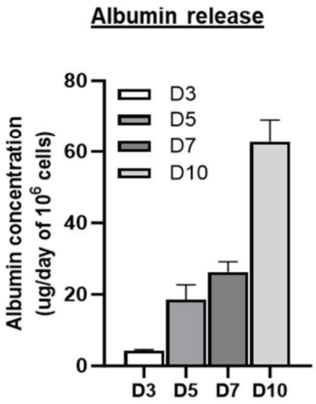


Figure S2. Albumin release of primary human hepatocyte grown on Curiochips across 10 days of culture.

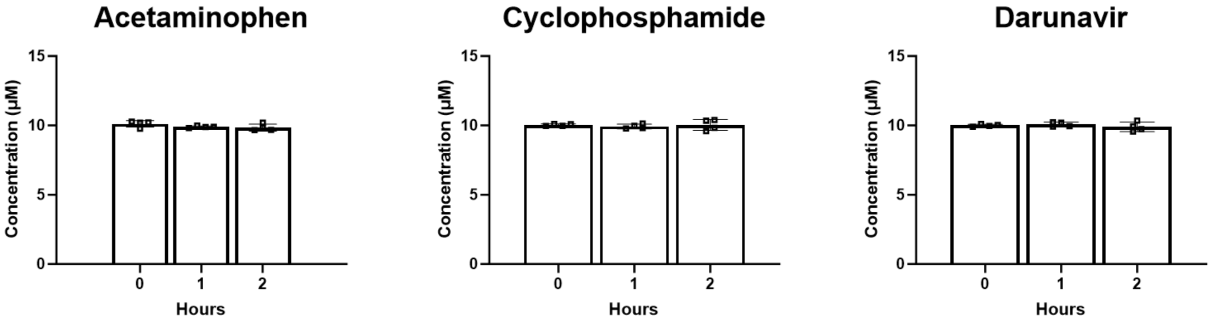


Figure S3. Acetaminophen, cyclophosphamide, and darunavir concentrations do not diminish after 2 hours on Curiochips, demonstrating no significant absorption of compounds by the microfluidic device.
